# Direct information estimation from cryo-EM Movies with CARYON

**DOI:** 10.1101/2020.11.25.398891

**Authors:** Kailash Ramlaul, Alister Burt, Natàlia de Martín Garrido, James T. MacDonald, Colin M. Palmer, Arjen Jakobi, Christopher H. S. Aylett

## Abstract

While cryo-EM with modern direct electron detectors has proven incredibly powerful, becoming a dominant technique in structural biology, the analysis of cryo-EM images is significantly complicated by their exceptionally low signal-to-noise ratio, limiting the accuracy of the parameterisation of the physical models required for successful classification and reconstruction.

Micrographs from modern direct electron detectors are typically collected as dose-fractionated multi-frame movies to allow the recording of separated individual electron impacts. These detectors improve electron detection and allow for both inter-frame motion correction, and dose-dependent image filtering, lessening the overall impact of effects deleterious to the recovery of high-resolution information.

In this study we measured the information content at each spatial frequency in cryo-EM movies as it accrues during the course of an exposure. We show that, as well as correction for motion and radiation damage, the use of the information within movies allows substantially improved direct estimation of the remaining key image parameters required for accurate 3D reconstruction: the image CTF and spectral SNR.

We are developing “CARYON” {insert contrived acronym here}, as a **LAFTER**-family filter for cryo-EM **movies** based upon such measurements. CARYON is intended to provide the best parameter estimation and filtration possible for a single complete, or large sub-section from a, movie micrograph without the use of a previously refined density. We demonstrate its utility in both single-particle and tomographic cryo-EM data processing.

## Introduction

### 1.0 Low-dose cryo-EM yields exceptionally noisy images, for which reliable parameter estimation remains problematic

Biological samples are highly radiation sensitive. Imaging of native-state biological samples is thus limited by the effects of radiation on the sample (Glaeser, 1971). Furthermore, biological samples are typically composed of predominantly light elements dissolved in water, and therefore the scattering contrast between sample and solvent is poor (Knapek & Dubochet, 1980). While electrons have been shown to yield the most scattering information of the types of ionising radiation typically used for biological imaging, only a very low electron dose can be applied during an image acquisition before sufficient damage occurs to render the high-resolution features of the sample unresolvable (Henderson, 1995). While electron scattering events occur according to a wave-form, they are detected as particles at a defined location. This results in “shot noise”, the loss of information due to the stochastic sampling of the scattering probability density. When these factors are coupled with those of radiation damage itself, the result is an incredibly low signal-to-noise ratio (SNR), which significantly hampers the extraction of usable information from electron micrographs.

### 1.1 “Movie” micrographs from direct electron detectors were developed to track individual electron impacts, but have been exploited for motion correction and dose-weighting

With the first movements towards direct electron detectors (McMullan *et al*, 2009), it became clear that minimising the impact footprint of each electron and assigning each impact equal weight would yield substantial improvements in the detective quantum efficiency, and therefore fractionation of the dose was performed to allow separation of electron impact events as they reached the detector, followed by individual impact detection and handling (Li *et al*, 2013; McMullan *et al*, 2014). Based upon this fractionation, further improvements that could be made were quickly put in place. Because microscopes are not entirely stable, exhibiting very low rates of drift, and more pertinently because samples exhibit further beam-induced movement due to charging as electron irradiation occurs (Russo & Henderson, 2018a, 2018b), motion correction was established (Brilot *et al*, 2012; Campbell *et al*, 2012; Li *et al*, 2013; Bai *et al*, 2013), to bring the information in frames into register. A slightly later advance in movie processing allowed for the down-weighting of image information as function of accumulated electron dose and therefore radiation damage (Grant & Grigorieff, 2015). “Dose weighting” is performed as a series of increasing B-factor filters based on the relative decay of information at different frequencies with dose, in order to maximise the contribution of the signal before it becomes degraded within each frequency band (Scheres, 2014; Grant & Grigorieff, 2015).

### 1.2 Despite early advances, comparisons between “frames” within movies have only recently been exploited in image correction

It is only recently that experimenters have begun to use the internal comparison available between frames within movies to perform estimation of further filters and parameters. This has principally been approached with deep-learning algorithms such as noise2noise (Lehtinen *et al*, 2018) in order to allow aggressive denoising of images by programs such as cryo-CARE, Warp and TOPAZ-denoise (Bepler *et al*, 2019). Despite their appealing visual aspect, aggressive real-space denoising approaches applied to cryo-EM micrographs are incompatible with high-resolution reconstruction. Instead, the low-SNR high frequency components of the image become dominated by higher-SNR phenomena at low frequencies, leading to loss of most of the high-resolution information. The shared information within frames should also be applicable to CTF estimation and classical Wiener filtering, though to our knowledge this has not yet been attempted. This could be because it has proven difficult to make accurate estimates of the signal strength in Fourier space from low SNR images.

### 1.3 Mixed real-space and Fourier measures allow improved parametrisation from low SNR “movie” images

The authors of this study, as well as Vilas and colleagues, and Pawel Penczek, have pioneered the use of “real-space measures” in SNR estimation, although to date this has been applied only to three-dimensional volumes recovered from the reconstruction process rather than two-dimensional images entering it. The LAFTER and SIDESPLITTER filters are based upon SNR estimation from band pass filtered real-space volumes (Ramlaul *et al*, 2019, 2020), while the more recently reported mFSC approach is similar (Penczek, 2020). All three methods allow substantially improved estimation of parameters from multiple independent low-SNR observations, and the general principle is applicable to any such problem.

### 1.4 We have developed CARYON to exploit the untapped information within “movie” images

In this manuscript we show that observing information accrual in multi-frame micrographs enables direct estimation of the Shannon information content within each frequency band (Shannon 1948). Because of the zeros of the CTF the contrast transfer function has nodes where no information is transferred (Zemlin, 1978), the information content of a cryo-EM image tends to zero at each such node, providing a relatively simple and robust target for CTF parameter optimisation. Our approach employs internal comparisons between the frames within a movie, and is readily generalised to equiphase bands when CTF correction is to be applied on a dose-fractionated cryo-EM “movie” micrograph (Zhang, 2016). Furthermore, under an assumption of smoothness, estimates of the SNR of each frequency band can be obtained, allowing direct, accurate Wiener-deconvolution of the images. {This approach will also allow direct local optimisation of and correction for the CTF when filtering tilted images, through the identification of nodes within real images of isolated equiphase bands, however this remains a work in progress at the time of writing}. For both cryo-electron tomography and compressed sensing applications, in which statistical estimation of the signal is otherwise exceptionally difficult, such a filter should prove very beneficial.

## Materials and Methods

### 2.0 Explicit assumptions underlying CARYON

Firstly, in order to perform information estimation using a LAFTER-like approach (Ramlaul *et al*, 2019), we must explicitly assume smoothness of the signal and noise spectra in reciprocal space. This is because we extract bands using a band-pass filter and then operate upon them in real-space, and any discontinuities will not be recovered correctly. We believe that this is a safe assumption because all biological macromolecules known to date are smooth at all resolutions currently accessible to cryo-EM. Secondly, in cases in which we are accounting for the CTF in two-dimensions (which is required for any accurate high-resolution filtering) we must extract bands along equiphase lines of the CTF. This implies that any significant quantity of astigmatism will lead to differential weighting of the same resolution band, which is highly undesirable. We therefore must assume that the astigmatism is negligible from the point of view of the resolution bands. This is unlikely to be a serious issue for the majority of applications as large amounts of astigmatism are considered unwise and typically avoided in cryo-EM data collection, however it is something that must considered when using CARYON. Thirdly, a further assumption is that the bandwidth over which the information is calculated is sufficiently smaller than the width of the CTF waveforms to allow them to be adequately represented. This issue can be avoided by up-sampling; however, it is worth noting that this is computationally intensive and has not been done in our reference implementation. Fourthly, we assume that the signal accrues linearly at each resolution with increasing exposure. This is a reasonable assumption, as the low-SNR nature of cryo-EM images means that saturation (diminishing returns) is highly unlikely except at the very lowest spatial frequencies. With large electron doses, however, over-filtering might occur due to loss of information due to radiation damage. Finally, during filtering we assume that the experimenter desires recovery of all information in the entire image. This will frequently not be the case because the objects of interest are small and low SNR in comparison to the larger higher-SNR features within the field of view. A high-pass filter, or under-correction of the lowest frequencies, will often be required for ease of interpretation (and will be provided in our reference implementation).

### 2.1 Information estimation

In Fourier space, the FSC and related measures are insufficient to allow accurate measurement of the information entropy (Supplementary Fig. 1). Real-space entropy estimation is much more accurate, which we believe to be in part for geometric reasons, due to the reduced dimensionality of the calculation, in part because the distribution is more tractable, as there are many extreme outliers in Fourier space, and in part for purely statistical reasons, due to the increased number of observations over which the summation is performed. We evaluated a series of different functions for information calculation; a K-nearest neighbours mutual information estimate, a histogram-type entropy estimate, and the explicit Gaussian assumption based on Pearson’s product moment correlation coefficient (Supplementary Fig. 1). All gave somewhat different results at extremely low resolution, however all converged quickly to essentially identical values at higher resolutions as was expected given the success of the assumptions of Gaussian distributions for signal and noise in Fourier space. All estimates also exhibited some systematic bias, however, especially at very high resolutions beyond the information limit. To identify bias, we calculated our mutual information measures based on successive differences between frames. This allows us to account for any persistent bias in measurements and derive an estimate of the reliability of the mutual information estimation based upon the variance of these figures. Once the CTF is defined, we also incorporate estimates of the variance at zero expected information by monitoring the nodes of the CTF.

### 2.2 CTF and inter-frame motion estimation

Given accurate raw estimates for the mutual information between half-images, it proved possible to estimate the CTF parameters by minimising the information recovery at the nodes of the CTF. We performed three-parameter simplex minimisation over the region up to the information-limit (estimated based upon the observation of a statistically significant difference in mutual information between the peak and node of the CTF). Motion estimation in x, y, and z, was optimised per-frame using the same model, with maximisation of the peak-node information difference as a target instead of minimisation of information at the node.

### 2.3 Filter estimation

We estimate the SNR from the mutual information of the bivariate Gaussian by reversal of the Shannon-Hartley channel capacity calculation (Shannon, 1948), accounting for the fact that the signal between the two independent images has been subjected to attenuation by the (additive white gaussian) channel twice yields the following expressions for the SNR in terms of Pearson’s correlation coefficient (r) and information (M):

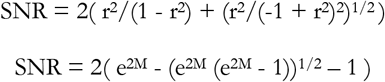

We corrected the SNR estimates according to the CTF (Fig. 2); crude estimates are made by division by the square of the CTF, and an improved estimate is then derived by Wiener-Kolmogorov estimation with a Gaussian kernel, using a maximum-likelihood approach to fit the kernel parameters. This has the effect of smoothing the estimated SNR spectrum, however we consider a slight smoothing to be desirable.

### 2.4 Reference implementation details

We note that the CARYON process is computationally intensive. We believe that further improvements and wider adoption will eventually necessitate the incorporation of elements of the CARYON process into single particle refinement programs and other pre-processing software, e.g. CTF estimation or motion correction. {CARYON is currently prototyped in Python using NumPy and SciPy, and code is available for CTF estimation, motion correction and filtering on request}. Before publication we intend to provide a reference implementation of CARYON as an optimised C99 program using FFTW3 for Fourier transformation (Frigo & Johnson, 2005), and the GNU scientific library implementation of simplex minimisation (Nelder & Mead, 1965)), to maximise speed and portability. CARYON operates on MRC mode 2 format image stacks (Cheng *et al*, 2015), i.e. C float or FORTRAN real. Source code for the CARYON reference implementation will be made available from the Imperial College Section for Structural Biology GitHub (github.com/StructuralBiology-ICLMedicine) under the GPL open source licence. CARYON can be run on any POSIX-compatible operating system, and will also eventually be made available in pre-compiled binary format for both Linux and Mac OS X as part of the CCP-EM suite (Burnley *et al*, 2017).

### 2.5 Data used for testing

For single particle analysis, we have used the Relion 3.1 test-dataset. For tomographic testing we used tilt series collected as dose-fractionated multi-frame micrographs of the E. coli mini-cell strain WM4196 (Burt *et al.* 2020).

## Results

### 3.0 The frequency-dependent information content of a movie can be effectively estimated from internal comparison

Estimates of mutual information in Fourier-space derived from the FSC, or attempted using nearest-neighbours, perform relatively well when large amounts of mutual information are present, but become extremely poor as the mutual information approaches zero (Supplementary Fig. 1). The estimation of mutual information from Fourier bands transformed to real-space, however, is reliable except at very low and very high frequencies (Fig. 1a; Supplementary Fig. 1). The accumulation of information can be traced at each frequency, and the resulting relationship is close to linear except at very low frequencies (Fig. 1b). These traces can be used to establish confidence in the estimated mutual information. Through correction for the effects of the CTF and the variance of the estimate, these mutual information estimates can be used to derive smooth SNR estimates suitable for use in Wiener deconvolution in the absence of excessive astigmatism (Fig. 2a).

**Figure 1:**
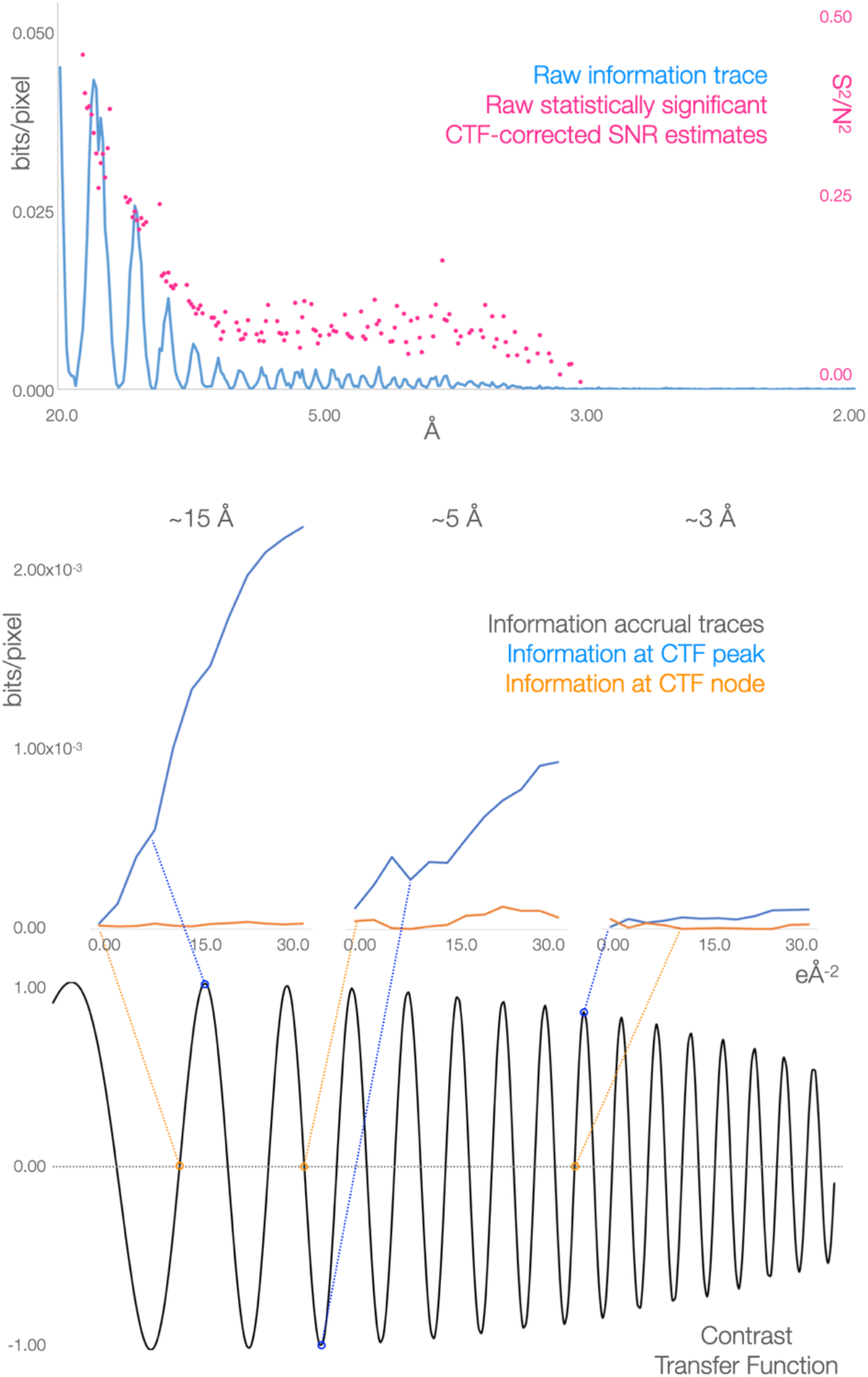
The information content of dose-fractionated movie images can be estimated reliably through the division of frames and the application of mixed real and Fourier space methods, allowing direct measurement of information accrual. [A] Curve showing the raw information estimated from half-images, and scatter showing the raw SNR estimated from the same. [B] Information accrual traces within a movie image at the indicated resolutions. The CTF is shown inset with the resolutions chosen highlighted.

**Figure 2:**
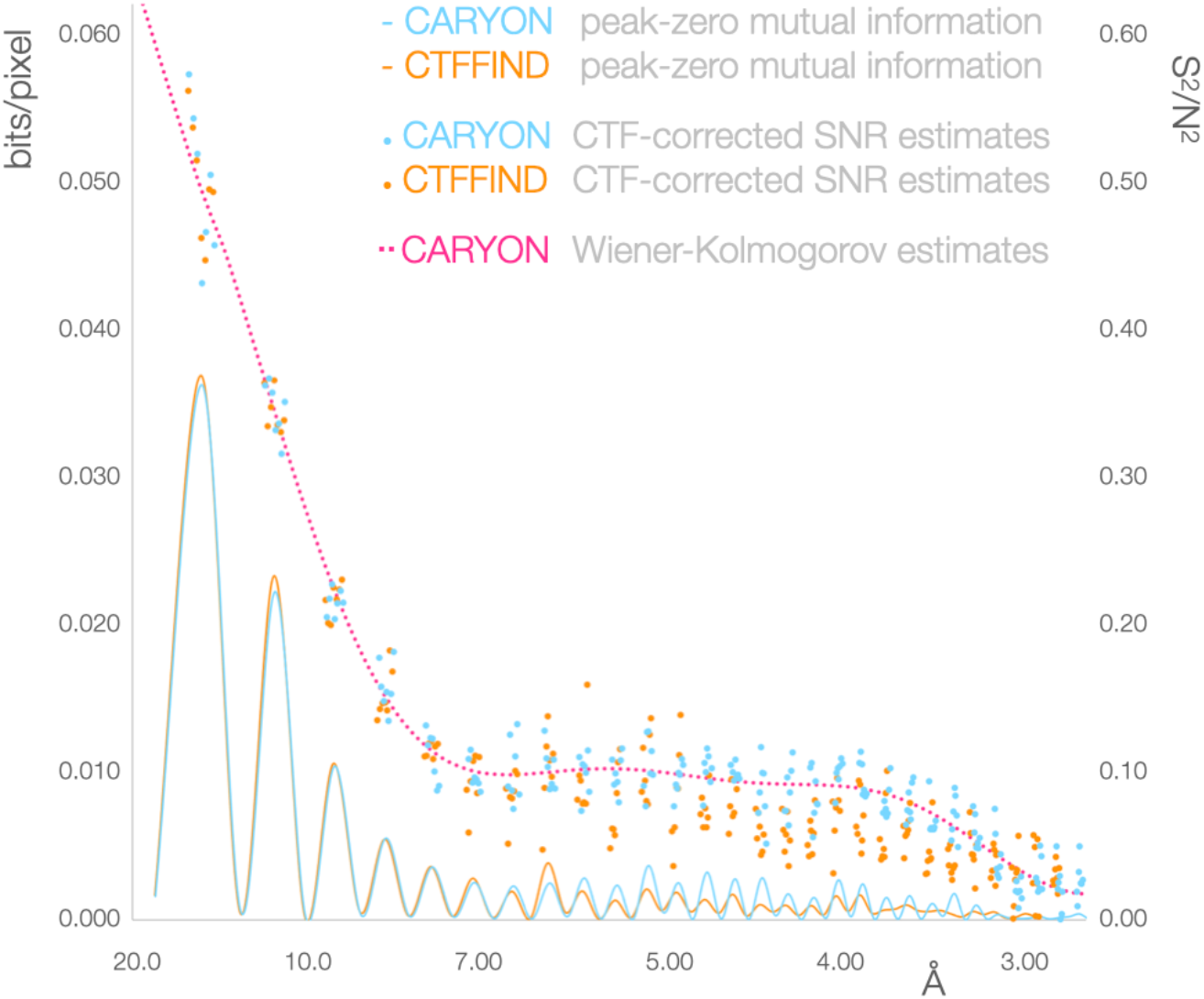
CTF and motion parameters can be improved by optimisation within movies using CARYON. [A] Direct comparison of radial information traces and raw SNR estimates from CTFFIND4 and CARYON, showing the improvement in recovery and SNR estimation through information estimation from frames as opposed to from the use of summed frames.

### 3.1 Minimisation of the raw information recovery at the nodes of the CTF allows efficient optimisation of the CTF parameters for a cryo-EM movie

Because the image transform is multiplied by the CTF, which has regular zeros, there should be no information along a series of equiphase lines within the CTF. By optimising the input CTF parameters to minimise the total amount of information recovered along these equiphase lines, an initial CTF estimate can be improved, and the resulting output fits are improved in comparison to the initial parameters (Fig. 2a). This approach requires at least several zeros to lie within the information limit, however this is expected to be the case for any interpretable cryo-EM movie image unless a phase-plate is in use.

### 3.2 Maximisation of the raw information difference between the peaks and nodes of the CTF below the information limit allows estimation of the x, y and z-offsets for frames within a cryo-EM movie

We corrected motion between individual frames by shifting them to maximise the information recovery between the peaks and nodes of the CTF. We do not see a substantial change in the estimated x- or y-shifts in comparison to the results of motioncor2 optimisation, which is expected given that the excluded regions are expected to lack signal, however by incorporating CTF information we can estimate shifts in z in addition to those in x and y, which is beneficial for per-frame filtering (Fig. 2b). {A similar approach – comparing halves from interleaved and initial versus final sets of frames – allows estimation of the information loss due to radiation damage, however these are no better than from standard dose-weighting at the time of writing}.

### 3.3 Images filtered with CARYON are superior for particle picking, and particles extracted from CARYON images can be classified more readily

Both CTF-deconvoluted and raw SNR filtered images can be generated with CARYON. Both types of CARYON output image provide improved visualisation of the features within the field of view and aid manual particle picking (Fig. 3a). When automated blob particle picking software (BOXER) is applied to original and CARYON-filtered images, the CARYON images demonstrate substantially improved particle picking (Fig. 3a). In contrast to neural network-based filters, CARYON filtered images can be used in single-particle refinement software because they retain information to high resolution. Refinements from CARYON SNR-filtered images reach the same, or close to the same, final resolution as unfiltered data. We believe that this will only have substantial benefits for single particle projects in extremely heterogenous cases, however, as refinement software can generate Wiener-deconvolution filters specific to the object of interest once refinement is convergent, which would be expected to perform better than filters estimated from the raw images alone. CARYON-filtered images demonstrate improved classification in 2D in comparison to raw input images (Fig. 3b). {Comparisons in 3D classification remain ongoing}.

**Figure 3:**
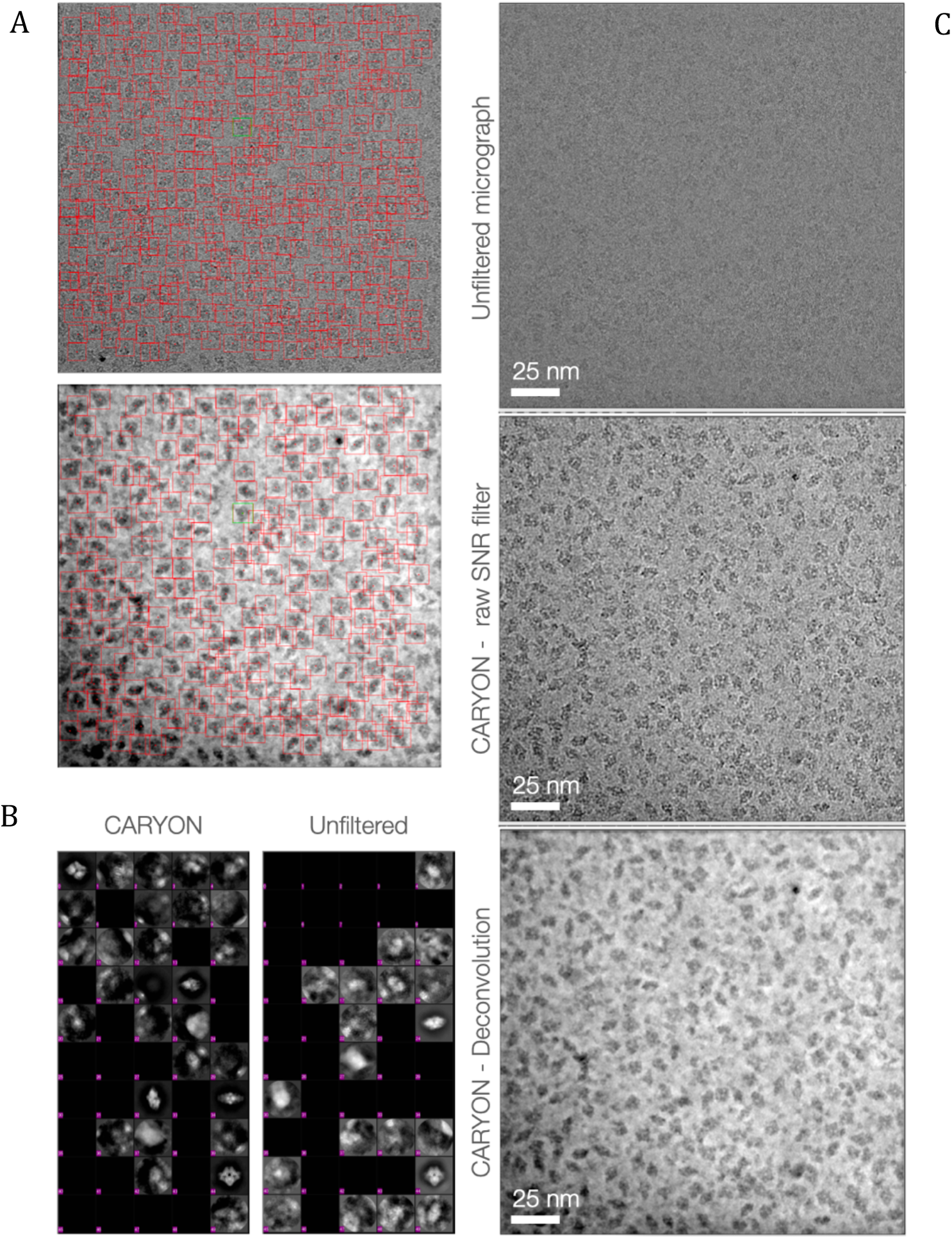
CARYON images improve particle picking and classification in single particle analysis. [A] Direct comparison of CARYON and input images during the particle selection process, showing improved contrast for manual particle picking and improved automatic particle selection. [B] Direct comparison of 2D classification from images with and without CARYON filtering – particle number is extremely low, leading to the collapse into two classes in the non-CARYON case. [C] Comparison of unfiltered, raw SNR, and deconvoluted images.

### 3.4 Images filtered according to CTF and SNR parameters determined with CARYON will be suitable for tomography

Images filtered according to SNR parameters generated in real-space are suitable for tomographic reconstruction {The SNR filtering used in the images shown is from an outdated version of the real-space SNR estimation processes we are developing - CARYON will allow the direct local correction of the CTF across a tilted image; this is still a work in progress at the time of writing, but will be incorporated before publication}, and tomograms prepared with such raw SNR filtering have a more pleasing visual aspect than corresponding reconstructions from raw images, more similar to the appearance of a deconvolved tomogram (Fig. 4a). Furthermore, template matching using such tomograms improves the appearance of the peaks in the correlation volume from template matching experiments (Fig. 4b). CARYON filtered tomograms, like single-particle images, will be suitable for sub-tomogram averaging, although we believe that this will once again only have substantial benefits in certain cases, as refinement software can generate Wiener-deconvolution filters specific to the object of interest from the current best estimate of the reconstructed density.

**Figure 4:**
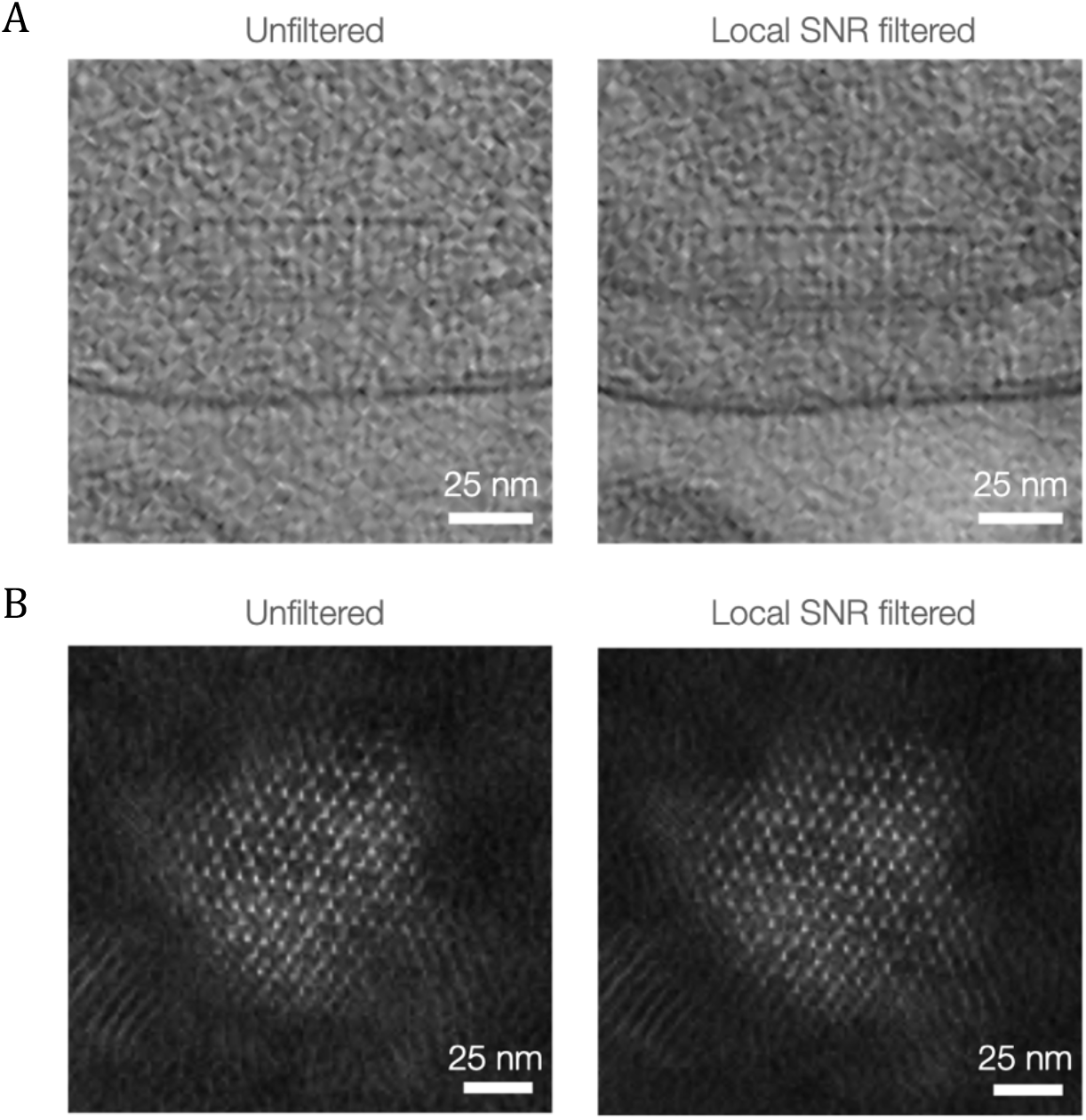
Real-space SNR filtered images improve template matching and tomographic reconstruction pipelines. [A] Comparison of sections from unfiltered and real-space SNR filtered tomograms. [B] Sub-tomogram template matching with and without real-space SNR filtration.

## Discussion

### 4.0 The CARYON methodology provides improved direct CTF, motion, and SNR estimates based on movies, which cannot replace model-derived estimates, but provide better starting points, and will substantially benefit cryo-EM applications in future

There is considerably more information available within a movie image than in the resulting average or sum. Current standard processing pathways attempt to make the best use of this information for the correction of drift and beam-induced motion (“motion correction”), as well as for differences in frequency-dependent degradation of the information over the course of the exposure (“dose-weighting”). To make the best use of the information available, in this study we show that movies can also be used for an internal comparison to improve the calculation of the parameters of the CTF and the band-dependent SNR.

In the case of CTF estimation, these calculations are computationally intensive in comparison to the methods currently in use to make these estimates, and therefore currently are likely to prove of greatest benefit in challenging situations where CTF estimation itself is problematic or where it is necessary to extend the CTF waveform to substantially higher resolution than the clearly identifiable Thon rings. This will typically be in low-dose situations such as individual tilt images in high-resolution tomography, when working with thick samples or for images from tilted data collections which have even lower SNR than usual. {CTF estimation from tilted images has been shown to be practicable and will be implemented before publication}.

In the case of SNR estimation and filtering, during single particle analysis and “sub-tomogram averaging”, which are now essentially similar processes from a computational perspective, the optimal filter is specific to the particle or sub-tomogram in question, and therefore calculation of the SNR or filter from an accurate, high-resolution model during the refinement process will remain the most powerful approach when such a model is readily achievable. Here we show that CARYON filtered images can be used for both 2D and 3D refinement, as the high-resolution components of the images remain accurately represented, even while the representation of the low-resolution components has been improved for visualisation and particle selection. In particular, this could mitigate the need for production of a separate tomogram for visualisation and particle picking purposes in cryo-ET. {CTF deconvolution across tilted images has also been shown to be practicable and will be implemented before publication}. Furthermore, we show improvement for 2D classification of problematic data, in the case shown due to low particle number. This has potential benefits in cases where the data display compositional or conformational heterogeneity, both of which require resolution before a high-resolution structure, and therefore appropriate filter, can be ascertained.

We believe that the principal application of CARYON, and future developments of these techniques, will lie in the improvement of tomographic image processing, where the CARYON filter will be beneficial as no suitable reliable volume for filter calculation is available, and in future developments towards low tilt number or “compressed sensing” approaches for biological cryo-EM. In such cases a reliable image deconvolution filter cannot be iteratively calculated from accurate high-resolution models as is currently the case for the averaging approaches, and approaches like those implemented within CARYON are essential. Furthermore, the use of the more-reliable real-space measures we propose here for CTF and SNR estimation, and of the local filtering we propose for Wiener-deconvolution of tilted images, will be of considerable benefit and can be directly incorporated into current single-particle or sub-tomographic refinement software where such accurate high-resolution models are available.

## Abbreviations

Cryo-EM: Cryogenic EM
CTF: Contrast transfer function
FSC: Fourier shell correlation
LAFTER: Local agreement filter for transmission EM reconstructions
SNR: Signal-to-noise ratio

## Acknowledgments

The authors would like to thank Takanori Nakane, Paul Simpson, Suhail Islam, and Rafael Ayala for their time and efforts during the development of the CARYON conceptual approach.

## Funding

This work was supported by the Wellcome Trust and the Royal Society through a Sir Henry Dale Fellowship (206212/Z/17/Z) to CHSA. AB was supported by a European Union Horizon 2020 research and innovation programme under grant agreement No 647784 to Irina Gutsche. CMP was supported by Medical Research Council funding (MR/N009614/1).

## Conflict of interest statement

The authors declare that they know of no conflicts of interest with respect to this work.

---

**Supplementary Figure 1:**
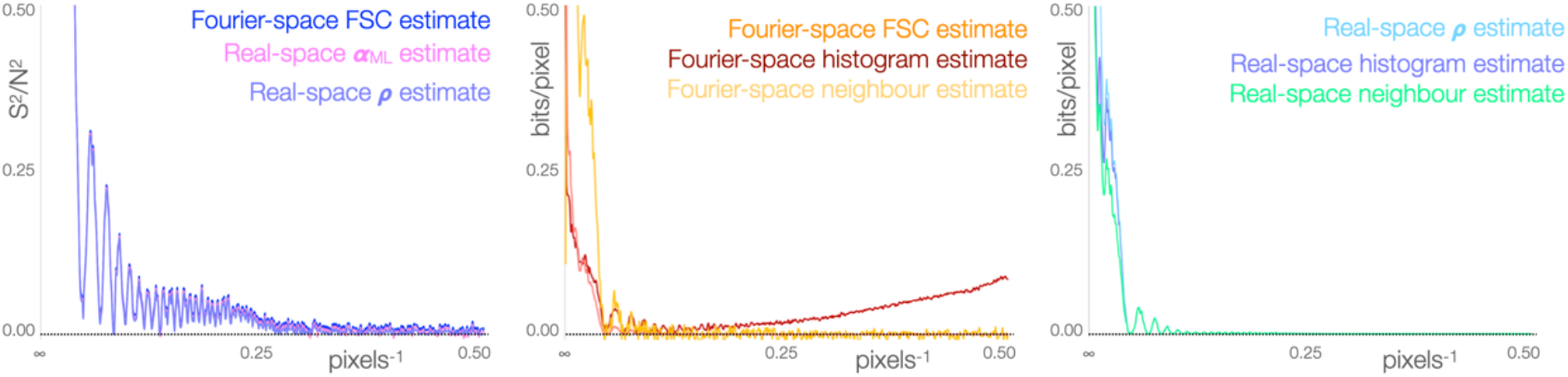
Mutual information estimates in real-space are consistent and reliable except at very low resolution, whereas those in Fourier space are unreliable and biased. [A] Comparison of raw SNR estimates derived from the FSC in Fourier space, the maximum-likelihood estimate in real-space, and the standard estimate from Pearson’s correlation coefficient in real-space, showing that the three estimates are convergent (they have been offset marginally in y to prevent curves from overlapping one another entirely), but noisy and relatively unsuitable, exhibiting residual variation to Nyquist. [B] Estimates of information per pixel in Fourier space made using the FSC and assuming the bivariate Gaussian information, a histogram binning approach, and the K-nearest neighbours’ approach, showing that all three estimates are noisy and exhibit residual variation to Nyquist, the FSC and histogram estimates are also highly inaccurate and unsuitable even after accounting for bias. [C] Estimates of information in real space made using Pearson’s correlation coefficient assuming the bivariate Gaussian information, a histogram binning approach, and the K-nearest neighbours’ approach. All three estimates are convergent, suitable, and reliable to Nyquist (see Fig. 1 for an expanded y-axis).

## Notes

### Competing Interest Statement

The authors have declared no competing interest.

## References

Bai XC, Fernandez IS, McMullan G & Scheres SHW (2013) Ribosome structures to near-atomic resolution from thirty thousand cryo-EM particles. Elife 2013:

Bepler T, Noble AJ & Berger B (2019) Topaz-Denoise: general deep denoising models for cryoEM. bioRxiv: 838920

Brilot AF, Chen JZ, Cheng A, Pan J, Harrison SC, Potter CS, Carragher B, Henderson R & Grigorieff N (2012) Beam-induced motion of vitrified specimen on holey carbon film. J. Struct. Biol. 177: 630–637

Burnley T, Palmer CM & Winn M (2017) Recent developments in the CCP-EM software suite. In Acta Crystallographica Section D: Structural Biology pp 469–477. International Union of Crystallography

Burt A, Cassidy CK, Ames P, Bacia-Verloop M, Baulard M, Huard K, Luthey-Schulten Z, Desfosses A, Stansfeld PJ, Margolin W, Parkinson JS, Gutsche I. (2020) Complete structure of the chemosensory array core signalling unit in an E. coli minicell strain. Nat Commun. 11: 743

Campbell MG, Cheng A, Brilot AF, Moeller A, Lyumkis D, Veesler D, Pan J, Harrison SC, Potter CS, Carragher B & Grigorieff N (2012) Movies of ice-embedded particles enhance resolution in electron cryo-microscopy. Structure 20: 1823–1828

Cheng A, Henderson R, Mastronarde D, Ludtke SJ, Schoenmakers RHM, Short J, Marabini R, Dallakyan S, Agard D & Winn M (2015) MRC2014: Extensions to the MRC format header for electron cryo-microscopy and tomography. J. Struct. Biol. 192: 146–150

Frigo M & Johnson SG (2005) The design and implementation of FFTW3. In Proceedings of the IEEE pp 216–231.

Glaeser RM (1971) Limitations to significant information in biological electron microscopy as a result of radiation damage. J. Ultrasructure Res. 36: 466–482

Grant T & Grigorieff N (2015) Measuring the optimal exposure for single particle cryo-EM using a 2.6 Å reconstruction of rotavirus VP6. Elife 4:

Henderson R (1995) The Potential and Limitations of Neutrons, Electrons and X-Rays for Atomic Resolution Microscopy of Unstained Biological Molecules. Q. Rev. Biophys. 28: 171–193

Knapek E & Dubochet J (1980) Beam damage to organic material is considerably reduced in cryo-electron microscopy. J. Mol. Biol. 141: 147–161

Lehtinen J, Munkberg J, Hasselgren J, Laine S, Karras T, Aittala M & Aila T (2018) Noise2Noise: Learning image restoration without clean data. ArXiv 7: 4620–4631

Li X, Mooney P, Zheng S, Booth CR, Braunfeld MB, Gubbens S, Agard DA & Cheng Y (2013) Electron counting and beam-induced motion correction enable near-atomic-resolution single-particle cryo-EM. Nat. Methods 10: 584–90

McMullan G, Chen S, Henderson R & Faruqi AR (2009) Detective quantum efficiency of electron area detectors in electron microscopy. Ultramicroscopy 109: 1126–1143

McMullan G, Faruqi AR, Clare D & Henderson R (2014) Comparison of optimal performance at 300keV of three direct electron detectors for use in low dose electron microscopy. Ultramicroscopy 147: 156–163

Nelder JA & Mead R (1965) A Simplex Method for Function Minimization. Comput. J. 7: 308–313

Penczek PA (2020) Reliable cryo-EM resolution estimation with modified Fourier shell correlation. IUCrJ 7:

Ramlaul K, Palmer CM & Aylett CHS (2019) A Local Agreement Filtering Algorithm for Transmission EM Reconstructions. J. Struct. Biol. 205: 30–40

Ramlaul K, Palmer CM, Nakane T & Aylett CHS (2020) Mitigating Local Over-fitting During Single Particle Reconstruction with SIDESPLITTER. J. Struct. Biol. 211: 107545

Russo CJ & Henderson R (2018a) Charge accumulation in electron cryomicroscopy. Ultramicroscopy 187: 43–49

Russo CJ & Henderson R (2018b) Microscopic charge fluctuations cause minimal contrast loss in cryoEM. Ultramicroscopy 187: 56–63

Scheres SH w (2014) Beam-induced motion correction for sub-megadalton cryo-EM particles. Elife 3: e03665

Shannon CE (1948) A Mathematical Theory of Communication. Bell Syst. Tech. journal. 27: 379–423

Tegunov D & Cramer P (2019) Real-time cryo-electron microscopy data preprocessing with Warp. Nat. Methods: 1–7

Vilas, J.L., Gómez-Blanco, J., Conesa, P., Melero, R., Miguel de la Rosa-Trevín, J., Otón, J., Cuenca, J., Marabini, R., Carazo, J.M., Vargas, J., Sorzano, C.O.S. (2018) MonoRes: Automatic and Accurate Estimation of Local Resolution for Electron Microscopy Maps. Structure. 26: 337–344

Zemlin F (1978) Image synthesis from electron micrographs taken at different defocus. Ultramicroscopy 3: 261–263

Zhang K (2016) Gctf: Real-time CTF determination and correction. J. Struct. Biol. 193: 1–12

